# Sequence-based prediction of vaccine targets for inducing T cell responses to SARS-CoV-2 utilizing the bioinformatics predictor RECON

**DOI:** 10.1101/2020.04.06.027805

**Authors:** Asaf Poran, Dewi Harjanto, Matthew Malloy, Michael S. Rooney, Lakshmi Srinivasan, Richard B. Gaynor

**Author notes:** These authors contributed equally (Asaf Poran, Dewi Harjanto, Matthew Malloy).

## Abstract

**Background:** The ongoing COVID-19 pandemic has created an urgency to identify novel vaccine targets for protective immunity against SARS-CoV-2. Consistent with observations for SARS-CoV, a closely related coronavirus responsible for the 2003 SARS outbreak, early reports identify a protective role for both humoral and cell-mediated immunity for SARS CoV-2.

**Methods:** In this study, we leveraged HLA-I and HLA-II T cell epitope prediction tools from RECON® (Real-time Epitope Computation for ONcology), our bioinformatic pipeline that was developed using proteomic profiling of individual HLA-I and HLA-II alleles to predict rules for peptide binding to a diverse set of such alleles. We applied these binding predictors to viral genomes from the *Coronaviridae* family, and specifically to identify SARS-CoV-2 T cell epitopes.

**Results:** To test the suitability of these tools to identify viral T cell epitopes, we first validated HLA-I and HLA-II predictions on *Coronaviridae* family epitopes deposited in the Virus Pathogen Database and Analysis Resource (ViPR) database. We then use our HLA-I and HLA-II predictors to identify 11,776 HLA-I and 7,991 HLA-II candidate binding peptides across all 12 open reading frames (ORFs) of SARS-CoV-2. This extensive list of identified candidate peptides is driven by the length of the ORFs and the significant number of HLA-I and HLA-II alleles that we are able to predict (74 and 83, respectively), providing over 99% coverage for the US, European and Asian populations, for both HLA-I and HLA-II. From our SARS-CoV-2 predicted peptide-HLA-I allele pairs, 368 pairs identically matched previously reported pairs in the ViPR database, originating from other forms of coronaviruses. 320 of these pairs (89.1%) had a positive MHC-binding assay result. This analysis reinforces the validity our predictions.

**Conclusions:** Using this bioinformatic platform, we identify multiple putative epitopes for CD4^+^ and CD8^+^ T cells whose HLA binding properties cover nearly the entire population and thus may be effective when included in prophylactic vaccines against SARS-CoV-2 to induce broad cellular immunity.

## Background

Coronaviruses are positive-sense single-stranded RNA viruses that have occasionally emerged from zoonotic sources to infect human populations (1). Most of the infections cause mild respiratory symptoms. However, some recent coronavirus infections have resulted in serious morbidity and mortality, including the severe acute respiratory syndrome coronavirus (SARS-CoV) (2–4), Middle East respiratory syndrome coronavirus (MERS-CoV) (5,6) and SARS-CoV-2, which is responsible for the current spreading of COVID-19. These three viruses all belong to the genus *Betacoronaviridae* (1). SARS-CoV was identified in South China in 2002 and its global spread led to 8096 cases and 774 deaths (7). The first case of MERS-CoV emerged in 2012 in Saudi Arabia, and since then a total of 2,494 cases and 858 associated deaths have been reported (6). In contrast to the more limited scope of these other coronavirus infections, SARS-CoV-2, which emerged in Wuhan, China at the end of December 2019, has resulted in 334,981 cases, including 14,652 deaths globally as of March 23, 2020 (8). The rapid spread of SARS-CoV-2 has resulted in the World Health Organization declaring a global pandemic. Thus, there is an urgent need for effective vaccines and antiviral treatments against SARS-CoV-2 to deal with this global pandemic.

The genome of SARS-CoV-2 spans 30 kilobases in length and encodes for 12 open-reading frames (ORFs), including four structural proteins. These structural proteins are the spike protein (S), the membrane protein (M), the envelope protein (E), and the nucleocapsid protein (N). In addition, there are over 20 non-structural proteins that account for all the proteins involved in the transcription and replication of the virus (9). All encoded proteins of the virus are potential candidates for developing vaccines to induce robust T cell immunity.

SARS-CoV and SARS-CoV-2 share 76% amino acid identity across the genome (10,11). This high degree of sequence similarity allows us to leverage the previous research on protective immune responses to SAR-CoV to aid in vaccine development for SARS-CoV-2. Both humoral and cellular immune responses have been shown to be important in host responses to SARS-CoV (12). Antibody responses generated against the S and the N proteins have shown to protect from SARS-CoV infection in mice and have been detected in SARS-CoV infected patients (13–16). However, these antibody responses detected against the S protein were short-lived and undetectable in patients six years post-recovery, suggesting that T cell responses may be involved in the long-term control of this virus (17). Indeed, significant changes in the total lymphocyte counts and T cell subset composition have been observed in patients with SARS-CoV; namely, levels of both B cells, and CD4^+^ and CD8^+^ T cells have been significantly reduced in these patients (18,19). Similarly, mice infected with SARS-CoV demonstrated that the severity of SARS correlated with the ability to develop a virus-specific T cell response (20,21).

Both CD4^+^ and CD8^+^ T cell responses have been detected in SARS-CoV-infected patients (12,22) as well as in SARS-CoV-2 (23). Notably, SARS-CoV-specific memory CD8^+^ T cells persisted up to 11 years post-infection in patients who recovered from SARS (24). Studies in mice have shown that SARS-CoV-specific memory CD8^+^ T cells provided protection against a lethal SARS-CoV infection in aged mice (21). In addition, adoptive transfer of effector CD4^+^ and CD8^+^ T cells to immunodeficient or young mice expedited virus clearance and improved clinical results (20). Immunization with SARS-CoV peptide-pulsed dendritic cells invigorated a T cell response, increasing the number of virus-specific CD8^+^ T cells enhancing both virus clearance and overall survival (25). These studies indicate an important role for T cell responses in controlling disease severity, virus clearance and conferring protective immunity to SARS-CoV infections. Given the homology between SARS-CoV and SARS-CoV-2, as well as emerging data on SARS-CoV-2 (23), cellular immune mechanisms might play a critical role in providing protection against SARS-CoV-2.

Here, we used T cell epitope prediction tools from the bioinformatic pipeline RECON® (Real-time Epitope Computation for ONcology) (26,27) to identify SARS-CoV-2 epitopes recognized by CD4^+^ and CD8^+^ T cells. RECON was trained on high-quality mono-allelic major histocompatibility complex (MHC) immunopeptidome data generated via mass spectrometry. The use of mass spectrometry allows for the high throughput, and relatively unbiased, collection of MHC binding data compared to traditional binding affinity assays, as well as the inclusion of important chaperone molecules. Additionally, the use of engineered mono-allelic cell lines avoids dependence on in-silico deconvolution techniques and allows for allele coverage to be expanded in a targeted manner.

With this approach, we generated data for 74 human leukocyte antigen (HLA)-I and 83 HLA-II alleles (Supplementary Tables 1 and 2). This mass spectrometry data enabled us to train neural network-based binding predictors that outperform the leading affinity-based predictors for both HLA-I (26) and HLA-II (27). Furthermore, we demonstrated in (27) that this improved binding prediction leads to improved immunogenicity prediction by validating on a data set of tetramer responses to a diverse collection of pathogens and allergens (28,29). Although RECON was originally developed to prioritize neoantigens for immunotherapy applications, it is agnostic to the source of peptide sequences evaluated and can be easily applied to peptides derived from pathogens as well. As validation to that end, the binding predictors from RECON were used to score *Coronaviridae* family peptides that had been assayed for T cell reactivity or MHC binding from the Virus Pathogen Resource (ViPR) database (30). The ViPR database integrates viral pathogen data from internally curated data, researcher submissions and data from various external sources. Our approach provides a significant improvement in both the breadth of predictions, and their validity, compared with a recent study that had a similar aim (31). We used the HLA-I and HLA-II binding predictors from RECON to predict the binding potential of peptide sequences from across the entire SARS-CoV-2 genome for a broad set of HLA-I and HLA-II alleles, covering the vast majority of USA, European, and Asian populations (Supplementary Table 3). Epitopes that were predicted to have a high likelihood of binding for multiple alleles could potentially be included in vaccines to stimulate CD4^+^ and CD8^+^ immune responses against this virus.

## METHODS

### Retrieval of *Coronaviridae* family T cell epitopes from ViPR

Experimentally determined epitopes for the *Coronaviridae* family for human hosts were retrieved from the Virus Pathogen Database and Analysis Resource (ViPR) (https://www.viprbrc.org/; accessed March 5 2020) (30). To build a validation dataset, both positives and negatives for T cell assays and MHC binding assays were obtained. Only assays associated with alleles identified with at least four-digit resolution and supported by RECON (Supplementary Table 1) were included for this analysis. Positive calls were prioritized – that is, if a given peptide-allele pair was assayed multiple times by a specific assay type and was determined to be positive in any single one of the assays, the peptide-allele pair was classified as positive. Specifically, the priority was given by the following order: Positive-High > Positive-Intermediate > Positive-Low > Positive > Negative (e.g., a peptide allele pairing that was assayed three times with the results Positive-High, Positive, and Negative were assigned a Positive-High result).

### Binding prediction for ViPR *Coronaviridae* family T cell epitopes

Peptide-HLA-I allele pairs in the ViPR validation dataset were scored using RECON’s HLA-I binding predictor, a neural network-based model trained on mass spectrometry data (26).

Similarly, peptide-HLA-II allele pairs in the ViPR validation dataset were scored using RECON’s HLA-II binding predictor, a recently published convolutional neural network-based model trained on mono-allelic mass spectrometry data (27). When applying the HLA-II binding predictor, we used the highest score for all 12-20mers within a given assay peptide. This is meant to account for the fact that the predictor is trained on ligands observed via mass spectrometry and may learn processing rules that are irrelevant for assays that do not incorporate processing and presentation.

### Retrieval of SARS-CoV-2 sequence

The GenBank reference sequence for SARS-CoV-2 (accession: NC_045512.2) was used for this study. All twelve annotated open-reading frames (orf1a, orf1b, S, ORF3a, E, M, ORF6, ORF7a, ORF7b, ORF8, N, and ORF10) were considered as sources of potential epitopes.

### Identification of HLA-I Epitopes

To identify candidate HLA-I epitopes, all possible 8-12mer peptide sequences from SARS-CoV-2 were scored with RECON’s HLA-I binding predictor. The HLA-I binding predictor was used to score SARS-CoV-2 peptides binding against 74 alleles, including 21 HLA-A alleles, 35 HLA-B alleles, and 18 HLA-C alleles. Peptide-allele pairs were assigned a percent rank by comparing their binding scores to those of 1,000,000 reference peptides for the same respective allele. Peptide-allele pairs that scored in the top 1% of the scores of these reference peptides were considered strong potential binders.

These top-ranking peptides were then prioritized based on expected USA population coverage (allele frequencies obtained from (32) – USA frequencies calculated as follows: 0.623*EUR0.133*AFA0.068*APA0.176*HIS), given all the alleles each peptide was expected to bind to (i.e., all the alleles for which the peptide scored in the top 1%). The estimate of population coverage for each peptide was calculated as

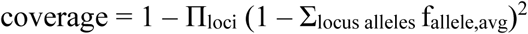

where f_allele,avg_ is the (unweighted) average allele frequency across the USA, European, and Asian Pacific Islander (API) populations and the cumulative product is taken across the three HLA-I loci: (HLA-A, HLA-B, and HLA-C).

The cumulative product itself represents the chance that an individual in the USA does not express any one of the contained alleles; hence, the complement describes the probability that at least one is present. The aim of using USA, European, and API allele frequencies is to cover a diverse population where allele frequency estimates are relatively reliable.

We then construct two ranked lists of HLA-I epitopes by coverage. The first ranks the epitopes by their absolute coverage, such that sequences predicted to bind similar collections of alleles would be ranked similarly (Supplementary Table 4). The second list, referred to as the “disjoint” list, is constructed in an iterative fashion where the sequence with the greatest coverage is selected first, and then the coverage for the remaining epitopes is updated to nullify contributions from any alleles that have already been selected (Supplementary Table 5). This second list was used to generate Figure 3A.

### Identification of HLA-II Epitopes

To identify HLA-II epitopes, we used RECON’s HLA-II binding predictor to score all 12-20mer sequences in the SARS-CoV-2 proteome to predict both binding potential and the likely binding core within each 12-20mer. Scoring was performed across all supported HLA-II alleles that comprise 46 HLA-DR alleles, 17 HLA-DP alleles, and 20 HLA-DQ alleles (Supplementary Table 2).

Peptide/allele pairs were assigned a percent rank by comparing their binding scores to those of 100,000 reference peptides. Pairs scoring in the top 1% were deemed likely to bind. Additionally, we define the “epitope” of a 12-20mer to be the predicted binding core within the sequence. As such, overlapping 12-20mers with the same predicted binding core for a given allele would constitute a single epitope. Table 1 shows counts of these epitopes.

**Table 1.** Summary of the HLA-I and HLA-II epitopes predicted across the 12 SARS-CoV-2 ORFs and their validation.

| ORFs | Length<br>(AA) | Peptide<br>HLA-I pair<br>count | Reported<br>in ViPR | Assay:<br>Negative | Assay:<br>Positive | Percent-<br>positive | Binding-<br>core and<br>HLA-II pair<br>count |
| --- | --- | --- | --- | --- | --- | --- | --- |
| envelope protein (E) | 75 | 556 | 34 | 3 | 31 | 91.2 | 29 |
| membrane<br>glycoprotein (M) | 222 | 1236 | 41 | 0 | 41 | 100.0 | 68 |
| nucleocapsid<br>phosphoprotein (N) | 419 | 1054 | 40 | 9 | 31 | 77.5 | 107 |
| orf1a polyprotein* | 4405 | 14* | 0 | 0 | 0 | NA | 0* |
| orf1ab polyprotein | 7096 | 28965 | 0 | 0 | 0 | NA | 2516 |
| ORF3a protein | 275 | 1408 | 127 | 11 | 116 | 91.3 | 94 |
| ORF6 protein | 61 | 322 | 0 | 0 | 0 | NA | 23 |
| ORF7a protein | 121 | 642 | 3 | 0 | 3 | 100.0 | 28 |
| ORF7b | 43 | 327 | 8 | 1 | 7 | 87.5 | 2 |
| ORF8 protein | 121 | 449 | 20 | 2 | 18 | 90.0 | 27 |
| ORF10 protein | 38 | 258 | 0 | 0 | 0 | NA | 4 |
| spike protein (S) | 1273 | 4686 | 95 | 14 | 81 | 85.3 | 437 |

\*peptides unique to orf1a (not found in orf1ab).

Additionally, we generated two lists of 25mers contained in SARS-CoV-2 protein sequences ranked by population coverage. To do this, we associated each 25mer with all subsequences that were likely binders and calculated the population coverage of the corresponding HLA-II alleles. Given a collection of alleles, we calculated the coverage as described in the previous section, the only difference being the cumulative product is taken across the following four HLA-II loci: HLA-DRB1, HLA-DRB3/4/5, HLA-DP, and HLA-DQ. HLA-II allele frequencies were obtained from (32) and Allele Frequency Net Database (33).

As with HLA-I, two sorted lists of predicted binding sequences were generated – one sorted on absolute coverage (Supplementary Table 6), and one sorted on disjoint coverage (Supplementary Table 7), which was used to generate Figure 3B and the observation that it would only require four 25mers to have predicted binders for >99.9% of the USA, European, and API populations.

### Comparison of predicted epitopes to the human proteome

8-12mer sequences (corresponding to predicted HLA-I epitopes), 9mer sequences (corresponding to predicted HLA-II binding cores), and 25mer sequences (corresponding to predicted HLA-II sequences that bound multiple alleles) from SARS-CoV-2 were compared against sub-sequences of the same length from the human proteome, using UCSC Genome Browser genes with hg19 annotation of the human genome and its protein coding transcripts (63,691 entries) (34). Exact matches were identified and flagged in Supplementary Table 4. No exact matches were found for the predicted HLA-II binding cores or 25mer sequences.

## RESULTS

### Validating RECON prediction for viral epitopes using ViPR

We first sought to validate the ability of our predictors to identify epitopes from genomes of the *Coronaviridae* family. Since SARS-CoV-2 only emerged recently, specific data on SARS-CoV-2 peptide MHC-binding and immunogenic epitopes are currently limited. However, other viruses from the *Coronaviridae* family have been studied thoroughly, specifically MERS-CoV and SARS-CoV. The latter has significant sequence homology to SARS-CoV-2 (35). We therefore sought to leverage previously tested epitopes from across the *Coronaviridae* family to validate our predictions of viral peptides, with special interest in peptide sequences that incidentally overlapped the novel SARS-CoV-2 virus. To that end, we used the publicly available ViPR database, which lists the results of T cell immunogenicity and MHC peptide-binding assays for both HLA-I and HLA-II alleles for viral pathogen epitopes. We used all assays of *Coronaviridae* family viruses with human hosts from ViPR as our validation dataset. Assays that did not have an associated four-digit HLA allele or were associated with an allele our models did not support were omitted (see Supplementary Tables 1 and 2 for a list of supported alleles).

For HLA-I, within the validation dataset there were a total of 4,445 unique peptide-HLA allele pairs that were assayed for MHC-binding, using variations of: 1) cellular MHC or purified MHC; 2) a direct or competitive assay; and 3) measured by fluorescence or radioactivity. Two additional peptide-MHC allele pairs were confirmed via X-ray crystallography. Depending on the study from which the data was collected, peptide-MHC allele pairs were either binarily defined in ViPR as “Negative” and “Positive” for binding, or with a more granular scale of positivity: Low, Intermediate, and High. We assigned peptide-MHC allele pairs with multiple measurements with the highest MHC-binding detected across the replicates (see Methods).

We then applied our HLA-I binding predictor from RECON to the peptide-MHC allele pairs in the validation dataset and compared the computed HLA-I percent ranks of these pairs with the reported MHC-binding assay results (Supplementary Table 8). A low percent rank value corresponds to high likelihood of binding (e.g., a peptide with a percent rank of 1% scores amongst the top 1% of the reference peptides). The percent ranks of peptide-MHC allele pairs that had a binary “Positive” result in the MHC-binding assay were significantly lower than pairs with a “Negative” result. Further, in the more granular positive results, stronger assay results (low < intermediate < high) were associated with increasingly lower percent ranks (Figure 1A). In addition, the two peptide-MHC alleles that were confirmed by X-ray crystallography were predicted as very likely binders, with low percent rank scores of 0.07% and 0.30%. These results demonstrate that our HLA-I binding predictor from RECON can reliably predict the HLA-I binding of peptides from proteins of the *Coronaviridae* family, to which SARS-CoV-2 belongs.

**Figure 1.**
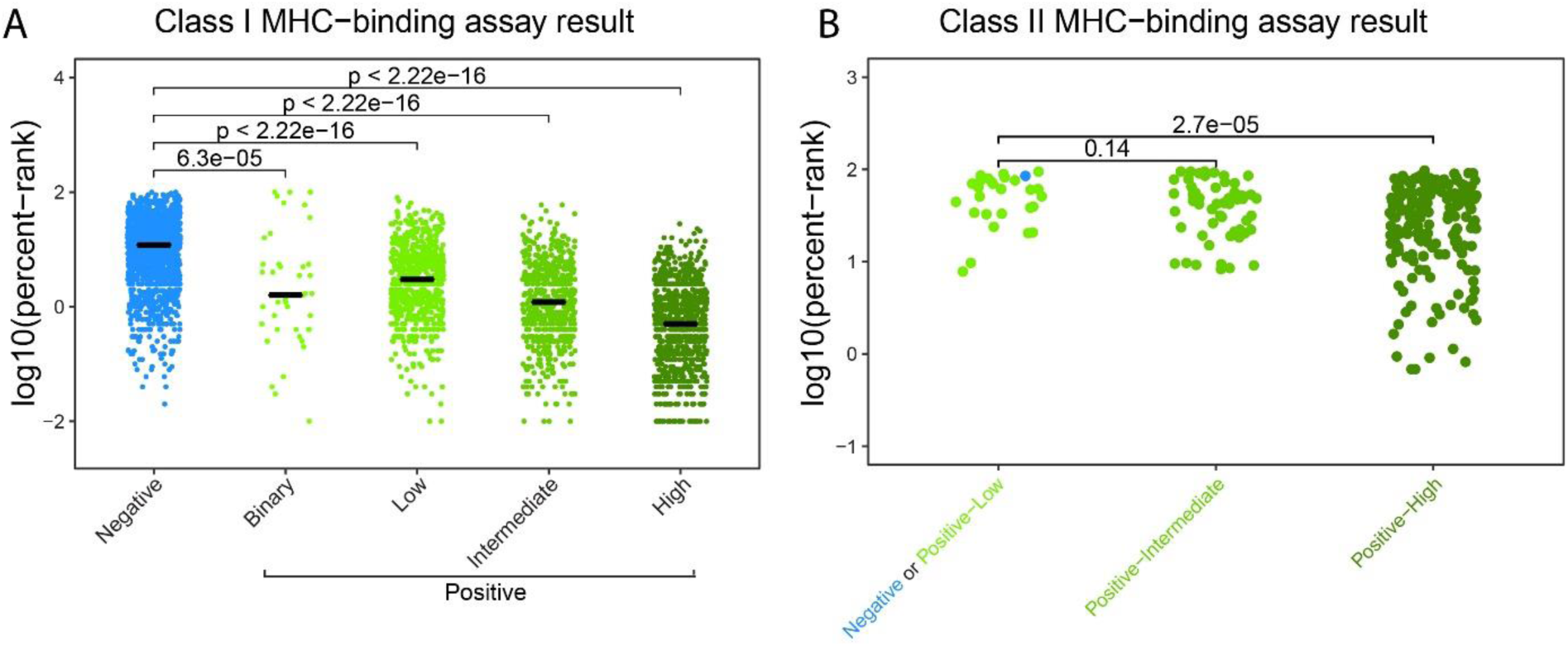
RECON binding predictors percent rank for both peptide- HLA-I and HLA-II binding pairs from ViPR correlate with their MHC-binding assay results. **A)** The log10(percent rank) of a predicted peptide-HLA-I allele pair, versus the ViPR reported MHC-binding assay result (either binary Negative/Positive or the scaled Negative/Positive-Low/Positive-Intermediate/Positive-High. In total, there were 4,445 peptide-HLA-I allele pairs. **B)** The log10(percent rank) of a predicted peptide-HLA-II allele pair, versus the ViPR reported MHC-binding assay result (Negative^+^Positive-Low/Positive-Intermediate/Positive-High). In total, there were 259 peptide-HLA-II allele pairs.

Assays of T cell reactivity (e.g., interferon-gamma ELISpots, tetramers), which are stricter measures for T cell immunogenicity to epitopes, were performed in significantly lower numbers compared with MHC-binding assays. For HLA-I, the overlap between peptide-MHC allele pairs for which we had a prediction (supported alleles) and pairs with a reported T cell assay consisted of only 32 pairs, of which 23 had a positive result. We did not detect differences in the percent ranks across the positive and negative groups, however sample sizes are extremely small (data not shown). In addition, for HLA-I epitopes, the validation dataset only contained T cell assay results for peptide-MHC allele pairs that had a positive result in a binding assay, suggesting a biased pool of epitopes selected for testing.

In addition to the identification of targets for CD8^+^ T cells, we have recently demonstrated and incorporated into RECON the unprecedented ability to predict HLA-II binders (27), allowing us to target CD4^+^ T cell responses which could be harnessed for SARS-CoV-2 vaccines. These CD4^+^ responses can potentially bolster both T cell immunity and enhance humoral immunity (36).

In a similar fashion to the HLA-I analysis, we scored all *Coronaviridae* family peptide-MHC allele pairs with supported HLA-II alleles in ViPR, using our HLA-II predictor (27) (Supplementary Table 9). There were 259 unique peptide-MHC allele pairs assayed by MHC-binding assays in the ViPR validation dataset for HLA-II. As before, we compared their percent rank with their reported ‘best’ (in the case of multiple measurements) MHC-binding assay result. This comparison could not be performed with the “Negative” pairs as an independent group since there was only one negative result in the validation dataset for HLA-II. The low negative counts may be due to under-reporting of negative assay results or biased selection of the peptides to be assayed. Therefore, we merged the “Negative” and “Positive-Low” groups into one group and compared their percent ranks with either the “Positive-Intermediate” or the “Positive-High” groups (Figure 1B). This analysis revealed a trend similar to that observed with HLA-I predictions, indicating that stronger MHC-binding assay results are associated with a lower predicted percent rank for HLA-II binders, as we expect for a robust predictor. Similar to the HLA-I T cell assays, there were too few recorded HLA-II T cell assays (31) in our validation dataset to determine percent rank differences between peptide-HLA II allele pairs testing positive and negative. Together, these findings further corroborate the validity of our epitope predictors, as peptide-MHC allele pairs with positive results in binding assays consistently have lower percent ranks (better scores) by both our HLA-I and HLA-II MHC-binding predictors.

### Epitope Prediction for SARS-CoV-2

We harnessed RECON’s HLA binding prediction ability to identify the peptides most relevant to the generation of SARS-CoV-2 T cell responses. We first performed the analysis for HLA-I peptide binding; we computed the likelihood of each peptide of lengths 8-12 amino-acid from the 12 SARS-CoV-2 ORFs to bind to any HLA-I allele in our database was computed. We then calculated the percent rank of each peptide-MHC allele pair by comparing their binding scores to those of a set of reference peptides, to generate a list of best-ranking peptide-MHC allele pairs (Figure 2A-C).

**Figure 2.**
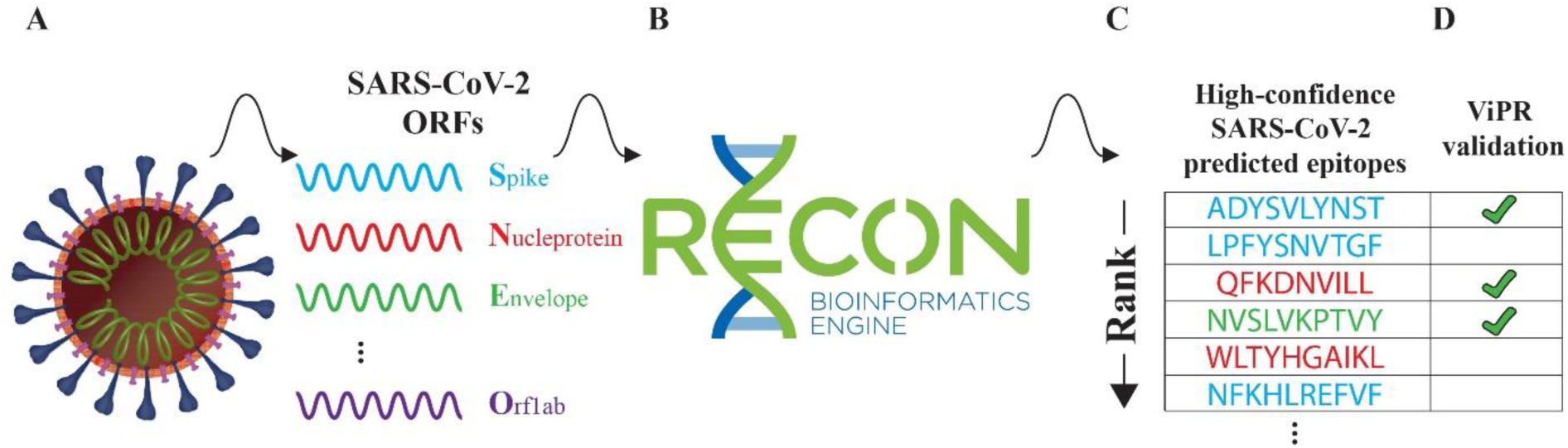
A schematic demonstrating our approach of applying the HLA binding predictors from RECON to identify SARS-CoV-2 T cell epitopes and validate using the ViPR database. **A)** A diagram of the SARS-CoV-2 virus, listing example proteins. **B)** Applying our HLA-I and HLA-II binding predictors from RECON to the 12 ORFs of SARS-CoV-2. **C)** Both HLA-I and HLA-II epitopes are ranked by their likelihood to bind a particular HLA allele (listed sequences were selected randomly and do not represent significant epitopes). **D)** Epitopes shared between SARS-CoV-2 and other coronaviruses which were previously assayed are used for validation.

We detected a total of 11,776 unique SARS-CoV-2 peptides that were predicted to bind at least one HLA-I allele with a percent rank score of 1% or lower (Supplementary Table 4). 14 of these peptides overlapped with a subsequence of the human proteome (see methods, Supplementary Table 4).

Unlike HLA-I, which has a closed binding groove that constrains bound peptide lengths to approximately 8 to 12 amino acids, peptides binding HLA-II have a wider length distribution (up to 30 amino acids or even longer) since the HLA-II binding groove is open at both ends. Peptides bind with a 9 amino acid subsequence (termed the binding core) occupying the HLA-II binding groove, with any flanking sequence overhanging the edges of the molecule. We consider a group of peptides that differ in the flanking regions but share a common binding core as a single epitope. Using the HLA-II predictor we identified 7,207 unique binding-cores that are predicted to bind at least one HLA-II allele with a percent rank score of 1% or lower. The number of high-quality peptide-MHC allele pairs we identify per SARS-CoV-2 gene is listed in Table 1. The majority of predicted peptide-MHC allele pairs are from orf1a and orf1ab, primarily driven by the length of these ORFs. In addition, orf1a and orf1ab have very similar sequences, with over 18,000 identical binding peptide-HLA-I allele pairs predicted for both ORFs. We therefore opted to exclude redundant predictions and only reported unique pairs (see * in Table 1). Similarly, all HLA-II predicted epitopes from orf1a were covered by those reported for orf1ab.

To test the validity of the SARS-CoV-2 predicted peptide-HLA pairs, we looked for identical peptide sequences in the *Coronaviridae* portion of the ViPR database (Figure 2D). A total of 368 HLA-I peptide-MHC allele pairs from SARS-CoV-2 had both a percent rank lower than 1% by our predictor and were found in the HLA-I MHC-binding validation dataset. Strikingly, of these HLA-I peptide-MHC allele pairs, 328 (89.1%) had a positive assay result. As a comparison, we also tested for overlap between epitopes predicted to have low likelihood of MHC-binding (percent rank 50% or higher) and the validation dataset. 37 peptide-MHC allele pairs overlapped between these sets, of which 36 (97.2%) had a negative assay result, as predicted. Further, we sought to determine whether our highly predicted SARS-CoV-2 peptide-HLA-I allele pairs (percent rank lower than 1%) would be validated by reported T cell assay results. Despite the significantly smaller number of peptide-MHC allele pairs that were tested for T cell reactivity in the validation dataset, 10 assayed pairs were also highly predicted by our HLA-I binding predictor. Nine out of these 10 (90%) predicted pairs had a positive result to the T cell assay. No low-scoring pairs (percent rank of 50% or above) were reported in the validation dataset. These findings demonstrate the validity of our prediction for peptide-HLA-I allele pairs for SARS-CoV-2 epitopes. Notably, while our algorithms are not trained on T cell reactivity data, and are aimed at peptide-MHC binding, for the few examples for which we had T cell reactivity assay results, we were able to show our highly-scoring peptide-MHC allele pairs are indeed immunogenic in the vast majority of cases.

For HLA-II peptide-MHC allele pairs, only a single HLA-II peptide-MHC allele pair had both a percent rank lower than 1% and was reported in the validation dataset; this single pair (from the envelope protein) had a “Positive-High” assay result.

### A small number of peptides predicted to bind multiple HLA-I and HLA-II alleles can provide broad population coverage

The concordance between the validation dataset and our highly predicted peptide-MHC allele pairs indicate that the HLA binding predictors from RECON significantly expand the list of predicted MHC binding peptides from the ORFs of SARS-CoV-2. We then sought to estimate the minimal number of HLA-I and HLA-II epitopes that would be required to provide coverage for the USA, European and Asian Pacific Islander populations based on the prevalence of MHC alleles in these populations (32). We found that a subset of the peptides was predicted to bind a broad set of either HLA-I or HLA-II alleles. We determined that a vaccine containing three of these HLA-I and four HLA-II sequences could provide >99.9% coverage for all of the USA, European, and Asian Pacific Islander populations, for HLA-I and HLA-II, respectively (Figure 3). Under the assumption that all peptide-MHC allele pairs for which a given peptide scores in the top 1% are indeed immunogenic, this finding could facilitate the design of a parsimonious, broadly effective vaccine.

**Figure 3.**
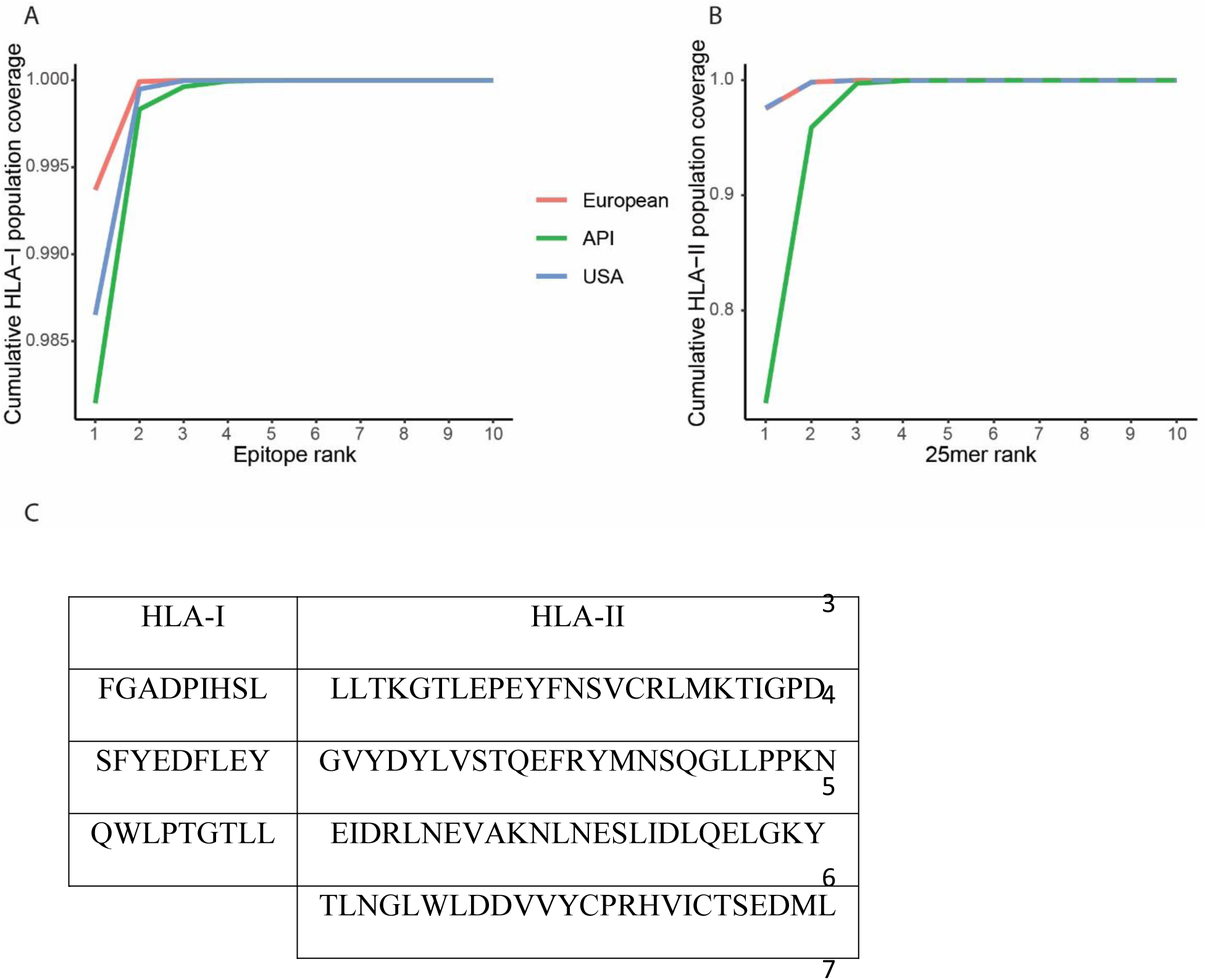
The USA, European and Asian Pacific Islander (API) populations could achieve >99% coverage with a small number of prioritized multi-allele binding epitopes. **A)** Cumulative HLA-I coverage for each population versus the number of included prioritized HLA-I epitopes. **B)** Cumulative HLA-II coverage for each population versus the number of included prioritized HLA-II 25mers. **C)** The amino-acid sequences predicted to cover over 99.9% of the populations in A and B.

## Discussion

In this work, we demonstrated the utility and validity of our HLA-I and HLA-II binding prediction algorithms to the *Coronaviridae* virus family, and specifically to SARS-CoV-2. By applying these algorithms to previously assayed peptide-MHC allele pairs in ViPR, we were able to show excellent concordance between our binding predictions and the results of the assays for both HLA-I and HLA-II epitopes. We leveraged the homology within the *Coronaviridae* family to demonstrate that an exceedingly high portion (~90%) of our high-ranking SARS-CoV-2 peptide-MHC allele pairs for which validation was available was indeed confirmed to bind the predicted MHC allele. Likewise, lowly-scoring peptide-MHC allele pairs derived from SARS-CoV-2 that had previously been assayed in ViPR were confirmed as non-binding. We therefore concluded that using RECON’s HLA binding predictors to predict T cell epitopes from the ORFs of SARS-CoV-2 provides a significantly expanded, novel set of high-quality vaccine targets for the virus. These sequences can be exploited for vaccines in various formats, including RNA or peptides.

This application of our prediction algorithms has clearly identified many candidate epitopes that can be included in a vaccine to induce cellular responses against this novel virus. Immune analysis of patients infected with either SARS-CoV or MERS-CoV has identified critical antiviral roles for these cellular responses. Viral specific CD8^+^ T cells can be cytotoxic and can kill virally infected cells to reduce disease severity. CD8^+^ T cells account for about 80% of the total inflammatory cells in the pulmonary interstitium of SARS-CoV infected patients and play a vital role in eliminating virally infected cells (12). In addition to having effector functions, CD4^+^ T cells can promote the production of virus-specific antibodies by activating T-dependent B cells. With respect to humoral immunity, antibodies are seen primarily to the S and N proteins. Although short lived, antibody responses are essential to control the persistent phase of CoV infection by preventing subsequent viral entry. We thus propose that a combination of B and T cell epitopes could provide long-lasting immunity from SARS-CoV-2 or mitigate the severity of disease when protection is partial.

The strength of our prediction is two-fold: first, we have validated predictors for both HLA-I and HLA-II binders, which potentially could be leveraged to induce both long-term CD4^+^ and CD8^+^ T cell immunity against the virus. Specifically, our HLA-II predictor, which has also been trained on a large set of mono-allelic mass spectrometry data, has been shown to significantly outperform previously published tools and is used here to identify high-quality CD4^+^ epitopes that may contribute to both cellular and humoral immunity (27) (Supplementary Table 6). Second, our expansive database of supported HLA-I and HLA-II alleles provides us with the ability to not only identify many peptide-MHC allele pairs, but to generate a narrow list of peptides with many potential HLA pairings that could be presented by the entire USA, European and Asian Pacific Islander populations. These advantages significantly improve upon previously published findings (31).

Our algorithms predict peptide-MHC binding, which is necessary but not sufficient to induce a T cell response. Therefore, further experimental work would be needed to refine the list of peptides to strictly immunogenic ones. However, with the breadth of the list we are able to provide, the likelihood of identifying many such epitopes is high. In addition, while the availability of confirmed T cell reactions to SARS2-CoV-2 epitopes is limited, nine out of 10 highly ranking peptide-HLA-I allele pairs that were previously assayed had a positive result in a T cell assay.

## Conclusions

In summary, our work provides the most robust set of both CD4^+^ and CD8^+^ T cells epitopes that are spanning the entire SARS-CoV-2 genome and binding a wide set of HLA-I and HLA-II alleles. Our predicted list of T cell epitopes serves as a resource for the scientific community to generate potent SARS-CoV-2 vaccine epitopes and generate long-lasting T cell immunity. These epitopes are predicted to bind MHC alleles covering over 99.9% of the USA, European and Asian Pacific Islander populations and could complement B cell epitopes that have been shown to be effective but provide short-lived immunity. This expansive data set allows us to identify peptides predicted to bind many alleles and to propose a small set of peptides that are predicted cover over 99.9% of USA, European, and Asian populations and induce broad CD8^+^ and CD4^+^ immunity.

## Supporting information

Function used for figure generation

## List of Abbreviations

S: Human Leukocyte Antigen
MERS-CoV: Middle East Respiratory Syndrome – Coronavirus
MHC: Major histocompatibility complex
RECON: Real-time Epitope Computation for ONcology
SARS-CoV: Severe Acute Respiratory Syndrome – Coronavirus
SARS-CoV-2: Severe Acute Respiratory Syndrome – Coronavirus – 2
USA: United States of America
ViPR: Virus Pathogen Resource
WHO: World Health Organization

## Declarations

### Ethics approval and consent to participate

Not applicable.

### Consent for publication

Not applicable.

### Availability of data and materials

All data generated or analyzed during this study are included in this published article and its supplementary information files.

### Authors’ contributions

A.P. – Conceptualization, Formal Analysis, Investigation, Visualization, Writing – Original Draft; D.H. – Conceptualization, Formal Analysis, Investigation, Visualization, Writing – Original Draft; M.M. – Conceptualization, Formal Analysis, Investigation, Visualization, Writing – Original Draft; M.S.R – Supervision, Writing – Review and Editing; L.S. – Supervision, conceptualization, Writing – Original Draft; R.B.G– Supervision, Funding Acquisition, Writing – Review and Editing; All authors read and approved the final manuscript.

## Acknowledgements

Not applicable.

## Additional files

<u>Supplementary Table 1: HLA-I Alleles Covered by Binding Predictors</u>

List of the 74 alleles covered by the HLA-I binding predictor.

Filename: SuppTable_1_RECON_classi_supported_alleles.csv

<u>Supplementary Table 2: HLA-II Alleles Covered by Binding Predictor</u>

List of the 83 alleles covered by the HLA-II binding predictor.

Filename: SuppTable_2_RECON_classii_supported_alleles.csv

<u>Supplementary Table 3: HLA-I and HLA-II Allele Population Frequencies</u>

HLA-I and HLA-II allele frequencies for USA, European (EUR), Asian Pacific Islander (API), African American (AFA), and Hispanic (HIS) populations. The USA population allele frequency is calculated as the following weighted average: 0.623*EUR+0.133*AFA+0.068*APA+0.176*HIS. For alleles where AFA and API population frequencies were not available, the USA population allele frequency values were set to match EUR. Missing API allele frequency values were imputed with 0 for our analyses.

Filename: SuppTable_3_classi_classii_allele_frequencies.xlsx

<u>Supplementary Table 4: Predicted HLA-I Binders Ranked by Population Coverage</u>

Table of all predicted HLA-I binders and their associated allele coverage. The table provides the peptide sequence, the SARS-CoV-2 protein(s) it is derived from, the HLA-I alleles it is predicted to bind to, the corresponding USA, European (EUR), and Asian Pacific Islander (API) population coverage, and a flag to indicate if the peptide sequence overlaps with any sequence in the human proteome.

Filename: SuppTable_4_classi_ranked_by_coverage.csv

<u>Supplementary Table 5: Broadly Binding HLA-I Peptides</u>

The 10 HLA-I predicted binders with the broadest cumulative allele coverage. The table provides the peptide sequence, its rank, the SARS-CoV-2 protein it is derived from, the alleles the peptide is predicted to bind to and the cumulative HLA-I coverage for USA, European (EUR), and Asian Pacific Islander (API) populations for all peptides up to this rank. See the *Identification of HLA-I Epitopes* section of the Methods for how coverage is calculated. Note that “surface glycoprotein” refers to the spike protein.

Filename: SuppTable_5_classi_best_cumulative_coverage.csv

<u>Supplementary Table 6: SARS-CoV-2 25mers Ranked by HLA-II Population Coverage</u>

Table of all SARS-CoV-2-derived 25mers containing at least 3 predicted HLA-II binders as subsequences. For each 25mer, the table provides the sequence, SARS-CoV-2 protein it is derived from, the alleles associated with the predicted binder subsequences, and their corresponding USA, European (EUR), and Asian Pacific Islander (API) population coverage.

Filename: SuppTable_6_covid_25mers_ranked_by_coverage_AVG.csv

<u>Supplementary Table 7: Broadly Binding HLA-II 25mers</u>

The 10 SARS-CoV-2-derived 25mers with the broadest cumulative predicted HLA-II allele coverage. For each 25mer, the table provides the rank, the peptide sequence, the SARS-CoV-2 protein it is derived from, the cumulative alleles that are covered by all 25mers up to this rank, and the associated USA, European (EUR), and Asian Pacific Islander (API) population coverage. Note that it is not the case that any of these 25mers, or their binding subsequences, are found as subsequences within the human proteome.

Filename: SuppTable_7_covid_25mers_best_cumulative_coverage_AVG.csv

<u>Supplementary Table 8: RECON binding prediction of ViPR HLA-I epitopes</u>

The peptide-HLA alleles pairs from the ViPR database which belong to the *Coronaviridae* family and have a human host had been scored using RECON’s HLA-I binding predictor. Alleles not reported in a four-digit format or not supported by RECON were excluded.

Filename: SuppTable_8_ViPR_classi_percent-rank.csv

<u>Supplementary Table 9: RECON binding prediction of ViPR HLA-II epitopes</u>

The peptide-HLA alleles pairs from the ViPR database which belong to the *Coronaviridae* family and have a human host had been scored using RECON’s HLA-II binding predictor. Filename: SuppTable_9_ViPR_classii_percent-rank.csv

## References

1. Cui J, Li F, Shi Z-L. Origin and evolution of pathogenic coronaviruses. Nat Rev Microbiol. 2019;17(3):181–92.

2. Gu J, Gong E, Zhang B, Zheng J, Gao Z, Zhong Y, et al. Multiple organ infection and the pathogenesis of SARS. J Exp Med. 2005 Aug 1;202(3):415–24.

3. Rota PA, Oberste MS, Monroe SS, Nix WA, Campagnoli R, Icenogle JP, et al. Characterization of a novel coronavirus associated with severe acute respiratory syndrome. Science. 2003 May 30;300(5624):1394–9.

4. Ksiazek TG, Erdman D, Goldsmith CS, Zaki SR, Peret T, Emery S, et al. A novel coronavirus associated with severe acute respiratory syndrome. N Engl J Med. 2003 May 15;348(20):1953–66.

5. WHO MERS-CoV Global Summary and Assessment of Risk. World Health Organization; 2018. https://www.who.int/csr/disease/coronavirus_infections/risk-assessment-august-2018.pdf

6. WHO | Middle East respiratory syndrome coronavirus (MERS-CoV). World Health Organization; [cited 2020 Mar 25]. http://www.who.int/emergencies/mers-cov/en/

7. WHO | Update 49 - SARS case fatality ratio, incubation period. World Health Organization; [cited 2020 Mar 25]. https://www.who.int/csr/sars/archive/2003_05_07a/en/

8. WHO | Coronavirus disease (COVID-19) Pandemic. [cited 2020 Mar 25]. https://www.who.int/emergencies/diseases/novel-coronavirus-2019

9. Chan JF-W, Kok K-H, Zhu Z, Chu H, To KK-W, Yuan S, et al. Genomic characterization of the 2019 novel human-pathogenic coronavirus isolated from a patient with atypical pneumonia after visiting Wuhan. Emerg Microbes Infect. 2020 Jan 28;9(1):221–36.

10. Zhou P, Yang X-L, Wang X-G, Hu B, Zhang L, Zhang W, et al. A pneumonia outbreak associated with a new coronavirus of probable bat origin. Nature. 2020;579(7798):270–3.

11. Lu R, Zhao X, Li J, Niu P, Yang B, Wu H, et al. Genomic characterisation and epidemiology of 2019 novel coronavirus: implications for virus origins and receptor binding. Lancet. 2020 22;395(10224):565–74.

12. Li G, Fan Y, Lai Y, Han T, Li Z, Zhou P, et al. Coronavirus infections and immune responses. J Med Virol. 2020;92(4):424–32.

13. Yang Z-Y, Kong W-P, Huang Y, Roberts A, Murphy BR, Subbarao K, et al. A DNA vaccine induces SARS coronavirus neutralization and protective immunity in mice. Nature. 2004 Apr 1;428(6982):561–4.

14. Graham RL, Becker MM, Eckerle LD, Bolles M, Denison MR, Baric RS. A live, impaired-fidelity coronavirus vaccine protects in an aged, immunocompromised mouse model of lethal disease. Nat Med. 2012 Dec;18(12):1820–6.

15. Wang J, Wen J, Li J, Yin J, Zhu Q, Wang H, et al. Assessment of immunoreactive synthetic peptides from the structural proteins of severe acute respiratory syndrome coronavirus. Clin Chem. 2003 Dec;49(12):1989–96.

16. Liu X, Shi Y, Li P, Li L, Yi Y, Ma Q, et al. Profile of antibodies to the nucleocapsid protein of the severe acute respiratory syndrome (SARS)-associated coronavirus in probable SARS patients. Clin Diagn Lab Immunol. 2004 Jan;11(1):227–8.

17. Tang F, Quan Y, Xin Z-T, Wrammert J, Ma M-J, Lv H, et al. Lack of Peripheral Memory B Cell Responses in Recovered Patients with Severe Acute Respiratory Syndrome: A Six-Year Follow-Up Study. The Journal of Immunology. 2011 Jun 15;186(12):7264–8.

18. Cui W, Fan Y, Wu W, Zhang F, Wang J, Ni A. Expression of lymphocytes and lymphocyte subsets in patients with severe acute respiratory syndrome. Clin Infect Dis. 2003 Sep 15;37(6):857–9.

19. Li T, Qiu Z, Zhang L, Han Y, He W, Liu Z, et al. Significant changes of peripheral T lymphocyte subsets in patients with severe acute respiratory syndrome. J Infect Dis. 2004 Feb 15;189(4):648–51.

20. Channappanavar R, Zhao J, Perlman S. T-cell-mediated immune response to respiratory coronaviruses. Immunol Res. 2014 Aug;59(0):118–28.

21. Channappanavar R, Fett C, Zhao J, Meyerholz DK, Perlman S. Virus-specific memory CD8 T cells provide substantial protection from lethal severe acute respiratory syndrome coronavirus infection. J Virol. 2014 Oct;88(19):11034–44.

22. Li CK, Wu H, Yan H, Ma S, Wang L, Zhang M, et al. T Cell Responses to Whole SARS Coronavirus in Humans. The Journal of Immunology. 2008 Oct 15;181(8):5490–500.

23. Thevarajan I, Nguyen THO, Koutsakos M, Druce J, Caly L, van de Sandt CE, et al. Breadth of concomitant immune responses prior to patient recovery: a case report of non-severe COVID-19. Nature Medicine. 2020 Mar 16;1–3.

24. Ng O-W, Chia A, Tan AT, Jadi RS, Leong HN, Bertoletti A, et al. Memory T cell responses targeting the SARS coronavirus persist up to 11 years post-infection. Vaccine. 2016 Apr 12;34(17):2008–14.

25. Zhao J, Zhao J, Perlman S. T Cell Responses Are Required for Protection from Clinical Disease and for Virus Clearance in Severe Acute Respiratory Syndrome Coronavirus-Infected Mice. J Virol. 2010 Sep;84(18):9318–25.

26. Abelin JG, Keskin DB, Sarkizova S, Hartigan CR, Zhang W, Sidney J, et al. Mass Spectrometry Profiling of HLA-Associated Peptidomes in Mono-allelic Cells Enables More Accurate Epitope Prediction. Immunity. 2017 21;46(2):315–26.

27. Abelin JG, Harjanto D, Malloy M, Suri P, Colson T, Goulding SP, et al. Defining HLA-II Ligand Processing and Binding Rules with Mass Spectrometry Enhances Cancer Epitope Prediction. Immunity. 2019 15;51(4):766–779.e17.

28. Archila LLD, Kwok WW. Tetramer-Guided Epitope Mapping: A Rapid Approach to Identify HLA-Restricted T-Cell Epitopes from Composite Allergens. Methods Mol Biol. 2017;1592:199–209.

29. Yang J, James EA, Huston L, Danke NA, Liu AW, Kwok WW. Multiplex mapping of CD4 T cell epitopes using class II tetramers. Clin Immunol. 2006 Jul;120(1):21–32.

30. Pickett BE, Sadat EL, Zhang Y, Noronha JM, Squires RB, Hunt V, et al. ViPR: an open bioinformatics database and analysis resource for virology research. Nucleic Acids Res. 2012 Jan;40(Database issue):D593–8.

31. Grifoni A, Sidney J, Zhang Y, Scheuermann RH, Peters B, Sette A. A Sequence Homology and Bioinformatic Approach Can Predict Candidate Targets for Immune Responses to SARS-CoV-2. Cell Host Microbe. 2020 Mar 12;

32. Maiers M, Gragert L, Klitz W. High-resolution HLA alleles and haplotypes in the United States population. Hum Immunol. 2007 Sep;68(9):779–88.

33. González-Galarza FF, Takeshita LYC, Santos EJM, Kempson F, Maia MHT, da Silva ALS, et al. Allele frequency net 2015 update: new features for HLA epitopes, KIR and disease and HLA adverse drug reaction associations. Nucleic Acids Res. 2015 Jan;43(Database issue):D784–788.

34. Kent WJ, Sugnet CW, Furey TS, Roskin KM, Pringle TH, Zahler AM, et al. The human genome browser at UCSC. Genome Res. 2002 Jun;12(6):996–1006.

35. Ahmed SF, Quadeer AA, McKay MR. Preliminary Identification of Potential Vaccine Targets for the COVID-19 Coronavirus (SARS-CoV-2) Based on SARS-CoV Immunological Studies. Viruses. 2020 25;12(3).

36. Zhao J, Zhao J, Mangalam AK, Channappanavar R, Fett C, Meyerholz DK, et al. Airway Memory CD4+ T Cells Mediate Protective Immunity against Emerging Respiratory Coronaviruses. Immunity. 2016 Jun 21;44(6):1379–91.

