## Supplementary material for "Sequence-based prediction of vaccine targets for inducing T cell responses to SARS-CoV-2 utilizing the bioinformatics predictor RECON": Function used for figure generation: Code_for_figures.txt

## Code for Manuscript Figures and Table 1

####Python
####
#### Calculate population fraction that will benefit as a function of peptides included
#### using cutoff of pct rank of 1 and below
####

import os
import sys
import pandas as pd
import numpy as np

import matplotlib
matplotlib.use('Agg')
import matplotlib.pyplot as plt
from matplotlib.backends.backend_pdf import PdfPages
from matplotlib.legend_handler import HandlerLine2D
from textwrap import wrap
from matplotlib.pyplot import cm

population = 'averaged' # averaged indicates using an aggregate of USA, API, and EUR population frequencies
omitted_peptides = [] # if want to ignore certain peptides - e.g. human proteome collisions
loci = ['A', 'B', 'C']

## load binding scores

df_scores = pd.read_csv('SARS-CoV-2.top2pct_neonmhc1.unique_peptide_alleles.scored.csv') # input: table with source protein(s), peptide, allele, neon_pctrnk 
df_scores = df_scores[df_scores['neon_pctrnk'] <= 1] # can further filter by %rank
peptides = sorted(set(df_scores['peptide']))
peptides = [i for i in peptides if i not in omitted_peptides]

## load allele frequencies

df = pd.read_table('collated_classi_allele_frequencies.txt') # input: table of allele frequencies
df['allele'] = df['allele'].apply(lambda x: 'HLA-' + x.replace('*',''))
df['averaged'] = df[['EUR','API', 'USA']].mean(axis=1)

## rank peptides by population coverage, quality of binding

def loadFreqs(df, population):
    allele2freq = dict(zip(df['allele'], df[population]))
missing_alleles = [i for i in set(df_scores['allele']) if i not in allele2freq] # patch frequencies for alleles with predictions that aren't in allele frequency table
for allele in missing_alleles:
    allele2freq[allele] = 0
return allele2freq

def normFreqs(locus, population):
    allele2freq = loadFreqs(df, population)
a2freq = {k: allele2freq[k] for k in [i for i in allele2freq.keys() if i.startswith(locus)]}
alleles = a2freq.keys()
freqs = a2freq.values()
numerator = 1/sum(freqs)
freqs = [numerator*i for i in freqs]
a2freq = dict(zip(alleles, freqs))
return a2freq

def calcCoverage(alleles, population):
    loci = ['HLA-A', 'HLA-B', 'HLA-C']
product = 1
for locus in loci:
    a2freq = normFreqs(locus, population)
locus_alleles = [i for i in alleles if locus in i]
summed_freqs = sum([a2freq[i] for i in locus_alleles])
product = product * (1 - summed_freqs)**2
coverage = 1 - product
return coverage

allele2freq = loadFreqs(df, population)

list_ = []
for peptide in peptides:
    alleles = ';'.join(sorted(df_scores[df_scores['peptide'] == peptide]['allele']))
allele_coverage = calcCoverage(alleles.split(';'), population)
scores = list(df_scores[df_scores['peptide'] == peptide]['neon_pctrnk']) # note scores aren't in same order as alleles
median_score = np.median(scores)
mean_score = np.mean(scores)
min_score = np.min(scores)
list_.append([peptide, alleles, allele_coverage, scores, median_score, mean_score, min_score])

df_flat = pd.DataFrame(list_, columns=['peptide', 'alleles', 'allele_coverage', 'neon_pctrnks', 'median_neon_pctrnks', 'mean_neon_pctrnks', 'min_neon_pctrnks'])
df_flat.sort_values(by=['median_neon_pctrnks'], inplace=True) # first sort by mean pct ranks to get best average scoring peptides - could also score by best ranking overall
if population != 'averaged':
    df_flat.drop_duplicates(subset='alleles', inplace=True) # drop peptides with duplicate allele representations

df_flat.sort_values(by=['allele_coverage'], inplace=True, ascending=False) # rank peptides by allele coverage
if population == 'averaged':
    df_flat['EUR_coverage'] = df_flat['alleles'].apply(lambda x: calcCoverage(x.split(';'), 'EUR_freq'))
df_flat['API_coverage'] = df_flat['alleles'].apply(lambda x: calcCoverage(x.split(';'), 'API_freq'))
df_flat['USA_coverage'] = df_flat['alleles'].apply(lambda x: calcCoverage(x.split(';'), 'global'))
df_flat['alleles_'] = df_flat['alleles'].apply(lambda x: ';'.join(sorted(x)))
df_flat = df_flat[['peptide', 'protein', 'alleles', 'alleles_', 'allele_coverage', 'USA_coverage', 'EUR_coverage', 'API_coverage', 'neon_pctrnks', 'median_neon_pctrnks', 'mean_neon_pctrnks', 'min_neon_pctrnks']]

df_flat['wt_collision'] = df_flat['peptide'].isin(omitted_peptides) # indicate if a peptide collides with the human proteome
df_flat.to_csv(population + '.peptide_pool_for_selection.csv', sep=',', index=False) # output: corresponds to supplemental table of all top-ranking peptides sorted by allele coverage

## rank peptides that maximize coverage

peptides = list(df_flat['peptide'])
peptide2alleles = dict(zip(df_flat['peptide'], df_flat['alleles']))

top_peptide = df_flat.iloc[0]['peptide']
banked_alleles = df_flat[df_flat['peptide'] == top_peptide]['alleles'].iloc[0].split(';')
banked_peptides = [top_peptide]

while len(banked_peptides) < 10:
    df_ = df_flat[~df_flat['peptide'].isin(banked_peptides)]
df_['new_alleles'] = df_['alleles'].apply(lambda x: [i for i in x.split(';') if i not in banked_alleles])
df_['new_coverage'] = df_['new_alleles'].apply(lambda x: 1 - np.prod([1-allele2freq[i] for i in x]))
df_.sort_values(by=['new_coverage'], inplace=True, ascending=False)
top_peptide = df_['peptide'].iloc[0]
banked_peptides.append(top_peptide)
banked_alleles = sorted(set(banked_alleles + df_flat[df_flat['peptide'] == top_peptide]['alleles'].iloc[0].split(';')))

list_ = []
if population == 'averaged':
    covered = []
for h, peptide in enumerate(banked_peptides):
    alleles = peptide2alleles[peptide].split(';')
covered = [i for i in alleles if i not in covered] + covered
data = [h+1, peptide, pep2orf[peptide], ';'.join(sorted(set(covered)))]
for population_ in ['averaged', 'EUR_freq', 'API_freq', 'global']:
    coverage = calcCoverage(covered, population_)
data.append(coverage)
list_.append(data)

df_cumulative = pd.DataFrame(list_, columns=['rank', 'peptide', 'protein', 'cumulative_alleles', 'averaged_AF', 'EUR_freq', 'API_freq', 'USA_freq'])
df_cumulative.to_csv(population + '.cumulative_freqs.disjoint_alleles.csv', sep=',', index=False) # output: corresponds to supplemental table of 10 peptides that maximize overall population coverage

## plot population coverage as a function of included peptides

def plotSaturation(peptides, fname):
    fig, ax = plt.subplots()
color=iter(cm.rainbow(np.linspace(0,1,5)))
pp = PdfPages(fname)
list_ = []
covered = []
for h, peptide in enumerate(peptides):
    alleles = peptide2alleles[peptide].split(';')
covered = [i for i in alleles if i not in covered] + covered
coverage = calcCoverage(covered, population)
list_.append([h+1, peptide, alleles, coverage])
df_coverage = pd.DataFrame(list_, columns=['rank', 'peptide', 'binder_alleles', 'population_coverage'])
df_coverage = df_coverage[:10]
c = next(color)
df_coverage.plot(y='population_coverage', x='rank', c=c, legend=False, ax=ax, alpha=0.5)
ax.set_title(population + ' allele frequencies') # , fontsize=6
ax.set_xlabel('Included peptides')
ax.set_ylabel('Population coverage')
plt.xticks(range(1,11), list(df_coverage['peptide']), rotation=90)
fig.tight_layout()
plt.savefig(pp,format='pdf',dpi=300)
plt.close(fig)
pp.close()

peptides = banked_peptides
plotSaturation(peptides, population + '.coverage_plots.disjoint_alleles.pdf') # output: corresponds to plot of population allele frequency in manuscript

##############
import numpy as np
import pandas as pd


def coverage_ranking(min_score=99, L_longmer=25, pop='AVG',
                     n_to_choose=12, min_peps=3):
    """
Return two dataframes of 25mers (for L_longmer=25):

df_selected - dataframe with n_to_choose rows, one per 25mer, ranked
by cumulative (pop) population coverage
df_abs - dataframe with one row per 25mer for all 25mers containing
at least min_peps predicted binders, sorted by absolute
(pop) population coverage. 

Binders are defined as any 12-20mers within the SARS-CoV-2 proteins
that are predicted to bind to an allele with model_score > min_score
Return list of 25mers within the nCoV proteins which cover the
largest fraction of the population based on the high-scoring
peptides they contain and the alleles they are predicted to bind to.
"""

# dummy function that returns all 12-20mers within SARS-CoV-2 proteins,
# and their predicted binding scores to covered alleles (model_score)
df_binders = get_all_covid_binders()
df_binders = df_binders.loc[df_binders['model_score'] >= min_score]
# this is a dummy function, would return dataframe with one row per
# 25mer within all the SARS-CoV-2 proteins
df_25mers = get_all_covid_25mers()

# dataframe of HLA-II population frequencies
df_freq = hla_freq()

# keep track of list of covered alleles and exclude them when calculating
# coverage in subsequent 25mer selections
excluded_hla = []
n_selected = 0
df_selected = pd.DataFrame()
while n_selected < n_to_choose:
    print('\n\nmaking selection %d of %d...' % ((n_selected+1), n_to_choose))
df_keep = df_binders.loc[~df_binders['allele'].isin(excluded_hla)]
df_list = df_keep[['peptide', 'allele']].groupby('peptide').aggregate(lambda tdf: tdf.unique().tolist())
df_list = df_list.reset_index()
pep_to_allele = pd.Series(df_list.allele.values, index=df_list.peptide).to_dict()
# associate each 25mer with peptides predicted to bind
binders = list(df_keep['peptide'].unique())
df_25mers['pep_list'] = \
df_25mers['peptide'].apply(lambda x: [y for y in binders if y in x])
# associate each 25mer with list of alleles the peptides bind to
df_25mers['allele_list'] = \
df_25mers['pep_list'].apply(lambda x: [pep_to_allele[y] for y in x])
df_25mers['alleles'] = \
df_25mers['allele_list'].apply(lambda x: [item for sublist in x
                                          for item in sublist])
df_25mers['alleles'] = df_25mers['alleles'].apply(lambda x: list(set(x)))
# calculate coverage of that allele list for various populations
df_25mers['coverage'] = \
df_25mers['alleles'].apply(lambda x: coverage(alleles=x,
                                              df_freq=df_freq,
                                              pop=pop))
df_25mers['USA_coverage'] = \
df_25mers['alleles'].apply(lambda x: coverage(alleles=x,
                                              df_freq=df_freq,
                                              pop='USA'))
df_25mers['EUR_coverage'] = \
df_25mers['alleles'].apply(lambda x: coverage(alleles=x,
                                              df_freq=df_freq,
                                              pop='EUR'))
df_25mers['API_coverage'] = \
df_25mers['alleles'].apply(lambda x: coverage(alleles=x,
                                              df_freq=df_freq,
                                              pop='API'))
print('\tzeroing out coverage for pep lists shorter than %d...' % min_peps)
df_25mers['npeps'] = df_25mers['pep_list'].apply(lambda x: len(x))
df_25mers.loc[df_25mers['npeps'] < min_peps, 'coverage'] = 0
df_25mers = df_25mers.sort_values('coverage', ascending=False)

# save absolute ranking
del df_25mers['allele_list']
if n_selected == 0:
    df_abs = df_25mers.copy()
df_abs['pep_list'] = df_abs['pep_list'].apply(lambda x: ';'.join(x))
df_abs['alleles'] = df_abs['alleles'].apply(lambda x: ';'.join(x))
if min_peps is not None:
    df_abs = df_abs.loc[df_abs['npeps'] >= min_peps]
df_abs = df_abs.loc[~df_abs['pep_list'].isnull()]
df_abs.to_csv('covid_%dmers_ranked_by_coverage_%s.csv' % (L_longmer, pop),
              sep=',', index=False)
df_25mers['all_alleles'] = df_25mers['alleles'].apply(lambda x: ';'.join(x))

# make next 25mer selection based on disjoint coverage
next_pick = df_25mers.iloc[0:1]
new_alleles = next_pick.iloc[0]['alleles']
next_pick['selection'] = n_selected + 1
# calculate cumulative coverage of all 25mers up to this point
next_pick['covered_alleles'] = next_pick['alleles'].apply(lambda x: x + excluded_hla)
next_pick['cumulative_coverage'] =\
next_pick['covered_alleles'].apply(lambda x: coverage(alleles=x, df_freq=df_freq,
                                                      pop=pop))
next_pick['USA_cumulative_coverage'] =\
next_pick['covered_alleles'].apply(lambda x: coverage(alleles=x, df_freq=df_freq,
                                                      pop='USA'))
next_pick['EUR_cumulative_coverage'] =\
next_pick['covered_alleles'].apply(lambda x: coverage(alleles=x, df_freq=df_freq,
                                                      pop='EUR'))
next_pick['API_cumulative_coverage'] =\
next_pick['covered_alleles'].apply(lambda x: coverage(alleles=x, df_freq=df_freq,
                                                      pop='API'))
df_selected = df_selected.append(next_pick)
n_selected += 1

excluded_hla += new_alleles


df_selected['pep_list'] = df_selected['pep_list'].apply(lambda x: ';'.join(x))
df_selected['alleles'] = df_selected['alleles'].apply(lambda x: ';'.join(x))

df_selected.rename(columns={'covered_alleles': 'cumulative_alleles',
    'alleles': 'additional_alleles',
    'all_alleles': 'alleles'},
    inplace=True)
df_selected['cumulative_alleles'] = \
df_selected['cumulative_alleles'].apply(lambda x: ';'.join(x))

del df_selected['coverage']
del df_selected['cumulative_coverage']
del df_selected['npeps']


return df_selected, df_abs


def hla_freq(pop_cols=None):
    """
Return HLA-II population frequencies for European (EUR),
Asian Pacific Islander (API), and United States (USA) populations,
as well as an average across the populations
"""

if pop_cols is None:
    pop_cols = ['USA', 'EUR', 'API']

hla_freq_file = # supplement hla-ii file
    df_hla = pd.read_csv(hla_freq_file, sep='\t')

df_dr = df_hla.loc[df_hla['locus'].isin(['DRB1', 'DRB345'])]

# impute missing allele frequencies with zeros
df_hla.fillna(0, inplace=True)

df_hla['AVG'] = df_hla[pop_cols].mean(axis=1)


return df_hla


def coverage(alleles=None, df_freq=None, pop='AVG'):
    """
Given a list of HLA-II alleles, calculate the population coverage of those
alleles where coverage is defined as the probability that a person
within that population has at least one of the alleles listed.
Function takes the population as an input as well as (optionally) the
population frequency table.

See the methods of the paper expression we use to calculate coverage.
"""

# if no alleles provide, pop_coverage = 0
if alleles == []:
    return 0

# account for possibility of duplicates in list
alleles = list(set(alleles))

# load frequency table if not provided
if df_freq is None:
    df_freq = hla_freq()

# dictionary {allele: frequency}
freq_dict = pd.Series(df_freq[pop].values,index=df_freq['allele']).to_dict()

# we are going to consider 4 loci, DRB1, DRB3/4/5, DP, DQ
dr1_alleles = [x for x in alleles if 'DRB1' in x]
bonus_alleles = [x for x in alleles if 'DR' in x and x not in dr1_alleles]
dp_alleles = [x for x in alleles if 'DP' in x]
dq_alleles = [x for x in alleles if 'DQ' in x]

hla_product = 1
if len(dr1_alleles) > 0:
    probs = sum([freq_dict[x] for x in dr1_alleles])
hla_product = hla_product * ((1 - probs) ** 2)
if len(bonus_alleles) > 0:
    probs = sum([freq_dict[x] for x in bonus_alleles])
hla_product = hla_product * ((1 - probs) ** 2)
if len(dp_alleles) > 0:
    probs = sum([freq_dict[x] for x in dp_alleles])
hla_product = hla_product * ((1 - probs) ** 2)
if len(dq_alleles) > 0:
    probs = sum([freq_dict[x] for x in dq_alleles])
hla_product = hla_product * ((1 - probs) ** 2)

pop_coverage = 1 - hla_product


return pop_coverage

########################################################
####R Code
## Figure 1 - ViPR Validation
library(tidyverse)
library(ggpubr)

class1_viper_wScores=read.csv("Supplementary Table 8.csv",header=T,stringsAsFactors = F)

comparisons_to_neg=list(c("Negative","Positive"),c("Negative","Positive-Low"),c("Negative","Positive-Intermediate"),c("Negative","Positive-High"))

class1_viper_wScores_MHCB=class1_viper_wScores %>% filter(Assay.Type.Category=="MHC Binding",!grepl("x-ray",Method))
class1_viper_wScores_MHCB$pmhc=paste(class1_viper_wScores_MHCB$Epitope.Sequence,class1_viper_wScores_MHCB$MHC.Allele.Name,sep = "_")
dups=unique(class1_viper_wScores_MHCB$pmhc)

reclass=NULL
for (i in dups){
    res_i=NULL
    dup_i=class1_viper_wScores_MHCB %>% filter(pmhc==i) %>% select(Assay.Result)
    tab_i=class1_viper_wScores_MHCB %>% filter(pmhc==i)
    if ("Positive-High" %in% dup_i$Assay.Result) {res_i="Positive-High"
    } else if ("Positive-Intermediate" %in% dup_i$Assay.Result) {res_i="Positive-Intermediate"
    } else if ("Positive-Low" %in% dup_i$Assay.Result) {res_i="Positive-Low"
    } else if ("Positive" %in% dup_i$Assay.Result) {res_i="Positive"
    } else if ("Negative" %in% dup_i$Assay.Result) {res_i="Negative"
    }
    reclass=rbind(reclass,data.frame(tab_i,correct=rep(res_i,nrow(dup_i))))
}

class1_viper_wScores_MHCB=reclass

class1_viper_wScores_MHCB_unique=class1_viper_wScores_MHCB %>% select(Epitope.Sequence,MHC.Allele.Name,correct,Neon.Percent.Rank)
class1_viper_wScores_MHCB_unique=unique(class1_viper_wScores_MHCB_unique)

class1_viper_wScores_MHCB_unique$correct=factor(class1_viper_wScores_MHCB_unique$correct,levels=c("Negative","Positive","Positive-Low","Positive-Intermediate","Positive-High"))

comparisons_to_neg=list(c("Negative","Positive"),c("Negative","Positive-Low"),c("Negative","Positive-Intermediate"),c("Negative","Positive-High"))

class1_viper_wScores_MHCB_unique %>%  
    ggplot(aes(x=correct,log10(Neon.Percent.Rank)))+geom_jitter(aes(color=correct),size=0.5,height = 0,width = 0.2)+stat_summary(fun.y = median, fun.ymin = median, fun.ymax = median,geom = "crossbar", width = 0.25)+
    stat_compare_means(method = "t.test",label="p.format",comparisons = comparisons_to_neg)+ylim(c(-2.5,4))+theme_classic()+theme_bw()+theme(panel.grid = element_blank(),axis.text.x = element_text(angle=45,hjust = 1))+xlab("MHC-binding assay result")+scale_color_manual(values = c("dodgerblue","chartreuse1","chartreuse2","chartreuse3","chartreuse4"))+guides(color="none")+ylab("log10(prediction percent-rank")

ggsave(filename = "Figure1A.pdf",width = 5,height = 4,useDingbats=FALSE)


##From the ViPR class 2 table MHC binding
class2=as_tibble(read.csv("Supplementary Table 9.csv",header=T,stringsAsFactors = F))
class2$Assay.Result=factor(class2$Assay.Result,levels=c("Negative","Positive","Positive-Low","Positive-Intermediate","Positive-High"))
class2$MHC.Allele.Name=gsub("\\*","",class2$MHC.Allele.Name)

class2_MHCB=class2 %>% filter(Assay.Type.Category=="MHC Binding",!grepl("x-ray",Method))
class2_MHCB$pmhc=paste(class2_MHCB$Epitope.Sequence,class2_MHCB$MHC.Allele.Name,sep = "_")
class2_MHCB_unique=class2_MHCB %>% select(Epitope.Sequence,MHC.Allele.Name,Assay.Result,Neon.Percent.Rank)
class2_MHCB_unique=unique(class2_MHCB_unique)

class2_MHCB_unique$Assay.Result2=class2_MHCB_unique$Assay.Result %>% as.character()
class2_MHCB_unique$Assay.Result2[class2_MHCB_unique$Assay.Result2 %in% c("Negative","Positive-Low")]="Negative-Positive-Low"
class2_MHCB_unique$Assay.Result2=factor(class2_MHCB_unique$Assay.Result2,levels=c("Negative-Positive-Low","Positive-Intermediate","Positive-High"))

comparisons_to_neg=list(c("Negative-Positive-Low","Positive-Intermediate"),c("Negative-Positive-Low","Positive-High"))

class2_MHCB_unique %>%  
    ggplot(aes(x=Assay.Result2,log10(100-Neon.Percent.Rank)))+geom_jitter(aes(color=Assay.Result),height = 0,width = 0.2)+
    stat_compare_means(method = "t.test",label="p.format",comparisons = comparisons_to_neg)+theme_classic()+theme_bw()+theme(panel.grid = element_blank(),axis.text.x = element_text(angle=45,hjust = 1))+xlab("MHC-binding assay result")+scale_color_manual(values = c("dodgerblue","chartreuse2","chartreuse3","chartreuse4"))+guides(color="none")+ylab("log10(percent-rank)")+ylim(c(-1,3))

ggsave(filename = "Figure1B.pdf",width = 5,height = 4,useDingbats=FALSE)


########################################################################
## Figure 3 - population coverage
## This code takes in the peptides ranked by their population coverage based on their predicted binding alleles, for either HLA-I (Supplementary Table 5) or HLA-II (Supplementary Table 7) 

library(tidyverse)
library(ggpubr)
library(reshape2)

# class i - columns "rank" and <API/EUR/USA>_freq
class1_cov=read.csv("Supplementary Table 5",header = T,stringsAsFactors = F)

class1_cov_forplot=melt(class1_cov[,c("rank","EUR_freq","API_freq","USA_freq")],id.vars = "rank")

ggplot(class1_cov_forplot,aes(x=rank,y=value,color=variable))+geom_line(size=1)+theme_bw()+theme_classic()+ylab("Cumulative HLA-I population coverage")+xlab("Epitope rank")+scale_x_continuous(breaks = 1:10)

ggsave(filename = "Figure3A.pdf",width = 5,height = 4,useDingbats=FALSE)

# class 2 - columns "selection" and <API/EUR/USA>_cumulative_coverage
class2_cov=read.table("Supplementary Table 7",header = T,stringsAsFactors = F,sep="\t")

class2_cov_forplot=melt(class2_cov[,c("selection","EUR_cumulative_coverage","API_cumulative_coverage","USA_cumulative_coverage")],id.vars = "selection")

class2_cov_forplot %>% filter(selection %in% 1:10) %>%
    ggplot(aes(x=selection,y=value,color=variable,linetype=variable))+geom_line(size=1)+theme_bw()+theme_classic()+ylab("Cumulative HLA-II population coverage")+xlab("25mer rank")+scale_x_continuous(breaks = 1:10)+scale_linetype_manual(values = c("solid","solid","dashed"))+guides(color="none",linetype="none")

ggsave(filename = "Figure3B.pdf",width = 5,height = 4,useDingbats=FALSE)


### Table 1 data
##From the ViPR class 1 table
scores_SARS2=read.csv("Supplementary Table 4.csv",header=T,stringsAsFactors = F)
scores_SARS2= scores_SARS2 %>% select(peptide, protein,alleles)

alleles_list=strsplit(scores_SARS2$alleles,split = ";")
proteins_list=strsplit(scores_SARS2$protein,split = ";")
proteins_list=sapply(proteins_list,tail,1)

scores_SARS2=data.frame(peptide=rep(scores_SARS2$peptide,sapply(alleles_list,length)),
                             allele=unlist(alleles_list),
                             protein=(rep(proteins_list,sapply(alleles_list,length))))

scores_SARS2$pmhc=paste(scores_SARS2$peptide,scores_SARS2$allele,sep="_")

hits_1per_counts=data.frame(table(scores_SARS2$protein))
names(hits_1per_counts)=c("protein","count")
hits_1per_counts$count[hits_1per_counts$protein=="orf1a polyprotein"]=length(setdiff(scores_SARS2$pmhc[scores_SARS2$protein=="orf1a polyprotein"],scores_SARS2$pmhc[scores_SARS2$protein=="orf1ab polyprotein"]))

hits_1per_counts$percent=round(100*hits_1per_counts$count/sum(hits_1per_counts$count),2)

class1=as_tibble(read.table("Supplementary Table 8.csv",header=T,sep=",",stringsAsFactors = F))
class1$Assay.Result=factor(class1$Assay.Result,levels=c("Negative","Positive","Positive-Low","Positive-Intermediate","Positive-High"))
class1$MHC.Allele.Name=gsub("\\*","",class1$MHC.Allele.Name)
class1_viper_wScores=merge(class1,scores_SARS2,by.x=c("Epitope.Sequence","MHC.Allele.Name"),by.y=c("peptide","allele"))

dups=unique(class1_viper_wScores$pmhc)

reclass=NULL
for (i in dups){
    res_i=NULL
    dup_i=class1_viper_wScores %>% filter(pmhc==i) %>% select(Assay.Result)
    tab_i=class1_viper_wScores %>% filter(pmhc==i)
    if ("Positive-High" %in% dup_i$Assay.Result) {res_i="Positive-High"
    } else if ("Positive-Intermediate" %in% dup_i$Assay.Result) {res_i="Positive-Intermediate"
    } else if ("Positive-Low" %in% dup_i$Assay.Result) {res_i="Positive-Low"
    } else if ("Positive" %in% dup_i$Assay.Result) {res_i="Positive"
    } else if ("Negative" %in% dup_i$Assay.Result) {res_i="Negative"
    }
    reclass=rbind(reclass,data.frame(tab_i,correct=rep(res_i,nrow(dup_i))))
}

class1_viper_wScores=reclass

class1_viper_wScores_unique=class1_viper_wScores %>% filter(Assay.Type.Category=="MHC Binding") %>% select(Epitope.Sequence,MHC.Allele.Name,correct,Neon.Percent.Rank,protein) 
class1_viper_wScores_unique=unique(class1_viper_wScores_unique)

class1_viper_wScores_unique$correct=factor(class1_viper_wScores_unique$correct,levels=c("Negative","Positive","Positive-Low","Positive-Intermediate","Positive-High"))
class1_viper_wScores_unique$pmhc=paste(class1_viper_wScores_unique$Epitope.Sequence,class1_viper_wScores_unique$MHC.Allele.Name,sep="_")
unique_to_1a=setdiff(scores_SARS2$pmhc[scores_SARS2$protein=="orf1a polyprotein"],scores_SARS2$pmhc[scores_SARS2$protein=="orf1ab polyprotein"])
unique_to_1a=nrow(class1_viper_wScores_unique %>% filter(protein=="orf1a polyprotein" & pmhc %in% unique_to_1a))

hits_1per_counts_SARS2=data.frame(table(class1_viper_wScores_unique$protein))
names(hits_1per_counts_SARS2)=c("protein","count")
hits_1per_counts_SARS2$count[hits_1per_counts_SARS2$protein=="orf1a polyprotein"]=unique_to_1a

hits_1per_counts_SARS2=cbind(hits_1per_counts_SARS2,matrix(table(class1_viper_wScores_unique$protein,class1_viper_wScores_unique$correct!="Negative"),ncol = 2))
names(hits_1per_counts_SARS2)=c("protein","count","Negative","Positive")
hits_1per_counts_SARS2$Negative[hits_1per_counts_SARS2$protein=="orf1a polyprotein"]=unique_to_1a
hits_1per_counts_SARS2$Positive[hits_1per_counts_SARS2$protein=="orf1a polyprotein"]=unique_to_1a

hits_1per_counts_SARS2$percent=round(100*hits_1per_counts_SARS2$Positive/hits_1per_counts_SARS2$count,2)

hits_1per_counts_full=merge(hits_1per_counts,hits_1per_counts_SARS2,by="protein",all=T)
hits_1per_counts_full[is.na(hits_1per_counts_full)]=0
names(hits_1per_counts_full)=c("protein","percent-rank 1 Epitope Count","fraction of all epitopes","Found in ViPR","Assay:Negative","Assay:Positive","percent positive")
